# Structural insights into xyloglucan recognition by an ABC transporter from a Gram-positive, thermophilic bacterium

**DOI:** 10.64898/2025.12.13.694138

**Authors:** Hansen Tjo, Virginia Jiang, Philip D. Jeffrey, Angela Zhu, A. James Link, Jerelle A. Joseph, Jonathan M. Conway

**Affiliations:** Department of Chemical and Biological Engineering, Princeton University, Princeton, NJ 08544, USA; Department of Molecular Biology, Princeton University, Princeton, NJ 08540, USA; Omenn-Darling Bioengineering Institute, Princeton University, Princeton, NJ 08544, USA; Andlinger Center for Energy and the Environment, Princeton University, Princeton, NJ 08544, USA; Department of Chemistry, Princeton University, Princeton, NJ 08540, USA; Princeton Institute for Computational Science and Engineering, Princeton University, Princeton, NJ 08544, USA; High Meadows Environmental Institute, Princeton University, Princeton, NJ 08544, USA

**Keywords:** xyloglucan, lignocellulose, carbohydrate-binding protein, ABC transporter, substrate-binding protein, *Anaerocellum bescii*, *Caldicellulosiruptor bescii*, thermophile, structural biology, crystallography

## Abstract

Xyloglucan (an α-1,6-xylosyl–substituted β-1,4-glucan) is a major hemicellulose of the primary cell wall of many plants and an important growth substrate for biomass-degrading bacteria in diverse ecological niches, including the gut microbiome and hot springs. In Gram-positive bacteria, xyloglucan is deconstructed into soluble oligosaccharides in the extracytoplasmic space before import by ATP-Binding Cassette (ABC) transporters, but the structural basis for this process remains poorly understood. Here, we identified an ABC transporter for xyloglucan uptake (Athe_2052-2054) in the Gram-positive, plant biomass-degrading thermophile *Anaerocellum bescii*, which is conserved across the *Anaerocellum* genus. We solved the apo crystal structure of its extracellular substrate-binding protein (SBP), Athe_2052, revealing a unique tertiary fold found only in a small subset of SBPs that bind complex oligosaccharides. This structure represents the first ABC SBP known to bind xyloglucan oligosaccharides. Biophysical analysis showed that while Athe_2052 binds unsubstituted β-glucan chains, recognition of xyloglucan side chains in the binding pocket markedly increases affinity (K_d_ = 14 nM) for xyloglucan heptasaccharide (XXXG), the principal oligosaccharide released during xyloglucan deconstruction. Molecular modeling revealed that xyloglucan heptasaccharide, owing to its branched substitutions, is bound in a distinct conformation compared to unsubstituted β-glucans. This represents a unique mode of xyloglucan recognition driven by α-linked side-chain interactions rather than β-glucan backbone recognition alone. Together, these findings provide the first structural basis for xyloglucan oligosaccharide recognition by an ABC transporter in Gram-positive bacteria.

## Introduction

Xyloglucan is a ubiquitous hemicellulose polysaccharide found in many dicot plants. It tethers the surface of cellulose microfibrils, conferring plant cell walls their rigidity, volume, and recalcitrance to enzymatic degradation (1,2). As a major component of the cell walls of cereals, fruits, and vegetables, xyloglucan can be found in many animal diets and is known to enrich the growth of gut microbiota that can degrade and utilize it in both human guts and cow rumen (1,3,4). Its abundance in plant biomass also makes it a valuable carbohydrate source for saprotrophs and lignocellulose degrading bacteria (5,6). Chemically, xyloglucan is composed of linear β-1,4-linked cello-oligosaccharides substituted with pendant α-1,6-linked xylose residues. Depending on the xyloglucan source, these xyloside side chains can be further substituted by galactose and fucose residues. Given xyloglucan’s biological importance, a molecular understanding of its utilization by microbes is of fundamental interest.

Gut symbionts prominent in complex plant polysaccharide degradation, such as those of the *Bacteroides* genus, have evolved a variety of proteins responsible for the binding, deconstruction, and utilization of xyloglucan (1,3). The anaerobic, spore-forming Gram-negative bacterium *Bacteroides ovatus* – found in the human colon – has garnered attention for its singular xyloglucan utilization locus (XyGUL), which encodes for a set of six enzymes encompassing eight glycoside hydrolase (GH) families that collectively degrade xyloglucan polysaccharide (1). The XyGUL also encodes for cell surface glycan-binding proteins (SGBPs) and a TonB-dependent transporter (TBDT) that moves xyloglucan oligosaccharides generated beyond the outer membrane (through the activity of endo-xyloglucanases GH5A, GH9A) into the periplasm, where they are further hydrolyzed into component monosaccharides by exo-glycosidases prior to cytoplasmic entry (1).

While the xyloglucan utilization system in *B. ovatus* – relying on cell surface glycan-binding platforms that bind very large oligosaccharides (degree of polymerization > 10) and a TBDT system – has been biochemically and structurally dissected, this is not the only known mechanism of xyloglucan uptake in microbes. Gram-positive bacteria such as *Ruminiclostridium cellulolyticum* have evolved a distinct mechanism whereby xyloglucan is deconstructed into soluble oligosaccharides directly in the extracytoplasmic space, followed by its direct uptake via ATP-Binding Cassette (ABC) transporters (5–7). Though this mechanism of xyloglucan utilization in Gram-positive bacteria has largely been studied biochemically and genetically, we do not yet have a structural basis for how these xyloglucan oligosaccharides are recognized by ABC transporter proteins.

ABC transporters that mediate carbohydrate uptake rely on extracellular substrate-binding proteins (SBPs) for substrate recognition. Unlike TonB-associated cell SGBPs, which coordinate large xyloglucan polysaccharides for deconstruction, ABC substrate-binding proteins recognize smaller, soluble oligosaccharides for cellular transport (8–10). The large expansion in unique SBP crystal structures in recent years (from 118 in 2010 to 764 in 2021) has necessitated their classification by structure and substrate preference (11–13). Contemporary frameworks currently group SBPs into eight distinct clusters A-H, differing in structural homology, size, sub-domain arrangement, and perhaps most importantly, substrate specificity, which spans siderophores and metals to complex carbohydrates and peptides (12,13). While AlphaFold2 provides highly accurate prediction of overall SBP folds and can configure ligand-binding pockets in some cases (14), for SBPs that recognize ligands absent or underrepresented in training data (e.g. branched oligosaccharides like XXXG), AlphaFold2 is not able to resolve an accurate ligand bound pose. Therefore, broadening the repertoire of experimentally determined SBP structures, together with characterization of their ligand interactions, is essential for advancing our understanding of protein-carbohydrate interactions with chemically complex carbohydrate ligands such as xyloglucan.

Structurally, SBPs are made up of two approximately symmetric globular α/β lobes joined by a flexible hinge region (8,11). Opposite this hinge region lies the binding pocket, where substrates are docked. A binding pocket can contain multiple sub-sites depending on its cognate substrate; each sub-site, comprising multiple residues, stabilizes one sugar monomer via hydrophobic and hydrophilic interactions (15,16). When a cognate substrate diffuses into an empty binding pocket, the two lobes pivot around the hinge region to close over the substrate, a process described as the “Venus-flytrap” system (11,15). This closed conformation is not only thermodynamically favored but also more thermally stable (16). Then, the SBP delivers the bound substrate to the ABC transmembrane domain (TMD) for translocation, after which it reverts back to its open, apo conformation.

Here, we explored the structure and function of the ABC transporter associated, xyloglucan-binding protein Athe_2052 from the Gram-positive bacterium *Anaerocellum bescii*. First, we dissected its ability to bind oligosaccharides, derived from tamarind seed xyloglucan digested by the native *A. bescii* secretome, and found that it binds xyloglucan heptasaccharide with the highest affinity. We also showed that Athe_2052 displays lower binding affinities for undecorated β-glucan chains, such as cellotriose and cellotetraose, that lack xylosyl substitutions. And as hemicellulose utilization is a core feature of members of the *Anaerocellum* genus, we found that Athe_2052 is highly conserved amongst members. Through X-ray crystallography, we solved the open, apo structure of Athe_2052 and discovered that it is structurally homologous to Cluster G SBPs (12,13). These SBPs are known for their unique tertiary fold and expansive binding pockets to accommodate large carbohydrate oligosaccharides; as only three other SBPs have been classified under Cluster G, this SBP subfamily is especially poorly characterized (12,13). Using molecular dynamics, we subsequently simulated the closed, holo form of Athe_2052 bound to its cognate ligand xyloglucan heptasaccharide. Our work offers new structural insights into xyloglucan utilization in Gram-positive bacteria, highlighting xylosyl side chain recognition by distinct SBP binding pocket subsites as a driving feature of xyloglucan binding specificity.

## Results

### The *Anaerocellum bescii* Secretome Degrades Tamarind Seed Xyloglucan into Oligosaccharides for Uptake by a Conserved ABC Transporter

The *A. bescii* secretome was isolated and used to digest tamarind seed xyloglucan to identify the released oligosaccharides. Digested xyloglucan was analyzed using liquid chromatography-mass spectrometry (LC-MS), showing that the *A. bescii* secretome degrades xyloglucan into the following oligosaccharides: xyloglucan heptasaccharide (XXXG), xyloglucan octasaccharide (XXLG), and xyloglucan nonasaccharide (XLLG) (Figure 1). No cellodextrin oligosaccharides or free xylose from further debranching were detected, suggesting that further enzymatic breakdown of xyloglucan oligosaccharides occurs intracellularly.

**Figure 1:**
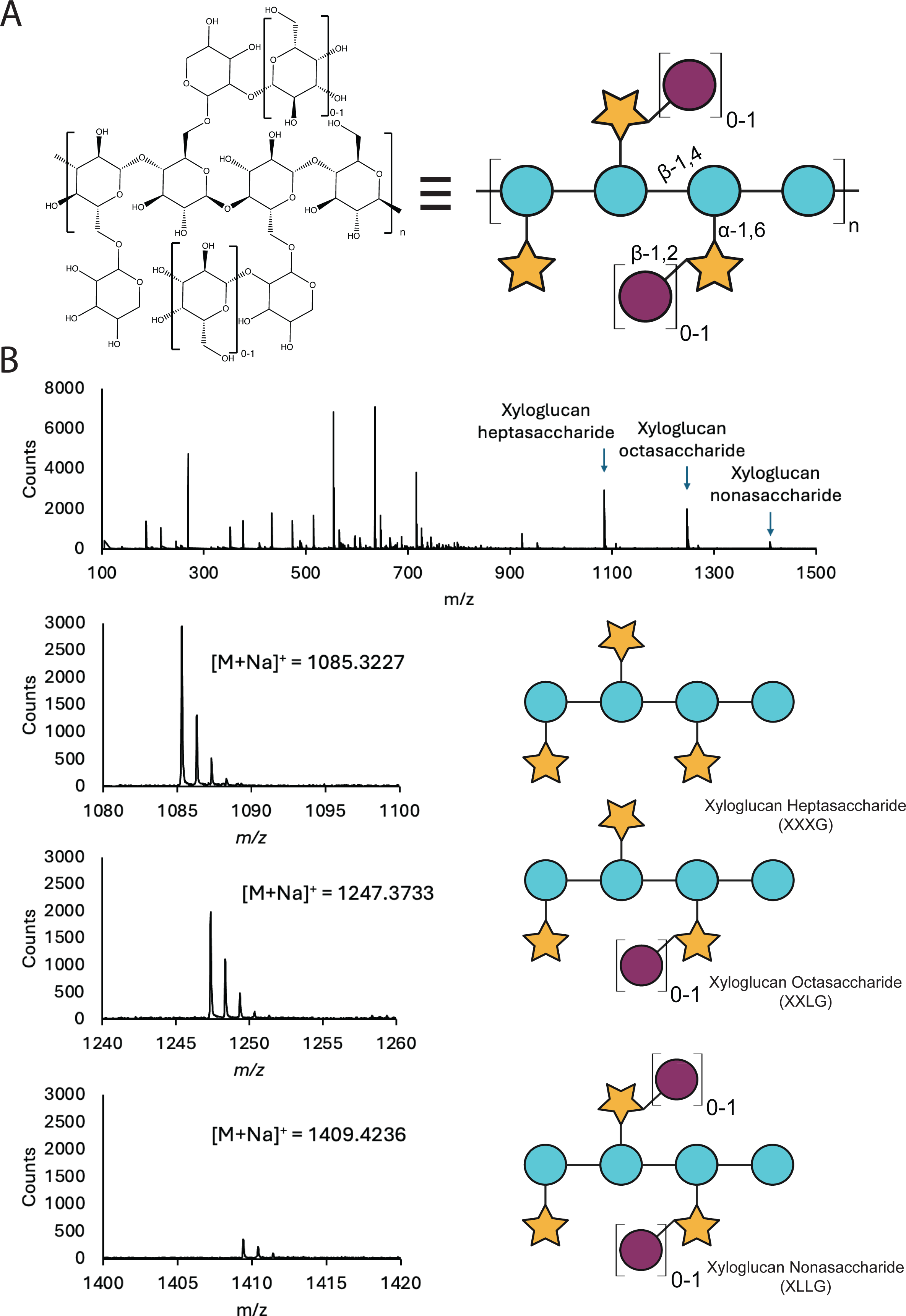
Xyloglucan hydrolysis by the *A. bescii* secretome yields XXXG, XXLG, and XLLG. **A)** Chemical structure of xyloglucan polysaccharide derived from tamarind seeds. **B)** Mass spectra showing the presence of xyloglucan heptasaccharide (XXXG), xyloglucan octasaccharide (XXLG), and xyloglucan nonasaccharide (XLLG) following enzymatic hydrolysis of tamarind seed xyloglucan. The expected monoisotopic *m/z* values are 1085.3379, 1247.3907, and 1409.4435, respectively. The retention time range for these mass spectra is 8.8 to 9.2 min. All species were identified as [M+Na]^+^ adducts. Glycan representations of the aforementioned xyloglucan oligosaccharides are included.

A previous transcriptomic study on *A. bescii* predicted ABC transporter locus Athe_2052-4 recognizes α-xylosides (17). The genomic context of this gene cluster suggests that it plays a role in xyloglucan oligosaccharide uptake (Figure 2). A co-localized, annotated α-xylosidase (encoded by *Athe_2057*) of GH family 31 likely represents the intracellular enzyme responsible for hydrolyzing the α-1,6-glycosidic bonds in xyloglucan oligosaccharides. *Athe_2053* and *Athe_2054* each encode for half of the transmembrane domain typical of the carbohydrate uptake transporter-1 (CUT1) Family of ABC transporters (Figure 2). CUT1 ABC transporters are known to possess a transmembrane domain encoded by two separate genes, and preferentially translocate larger, oligomeric substrates with the help of their extracytoplasmic SBP. Though no ATPase is located in proximity of the Athe_2052-4 locus, transport-coupled ATP hydrolysis in *A. bescii* CUT1 transporters is known to be reliant on the promiscuous ATPase MsmK, encoded by *Athe_1803* (18).

**Figure 2:**
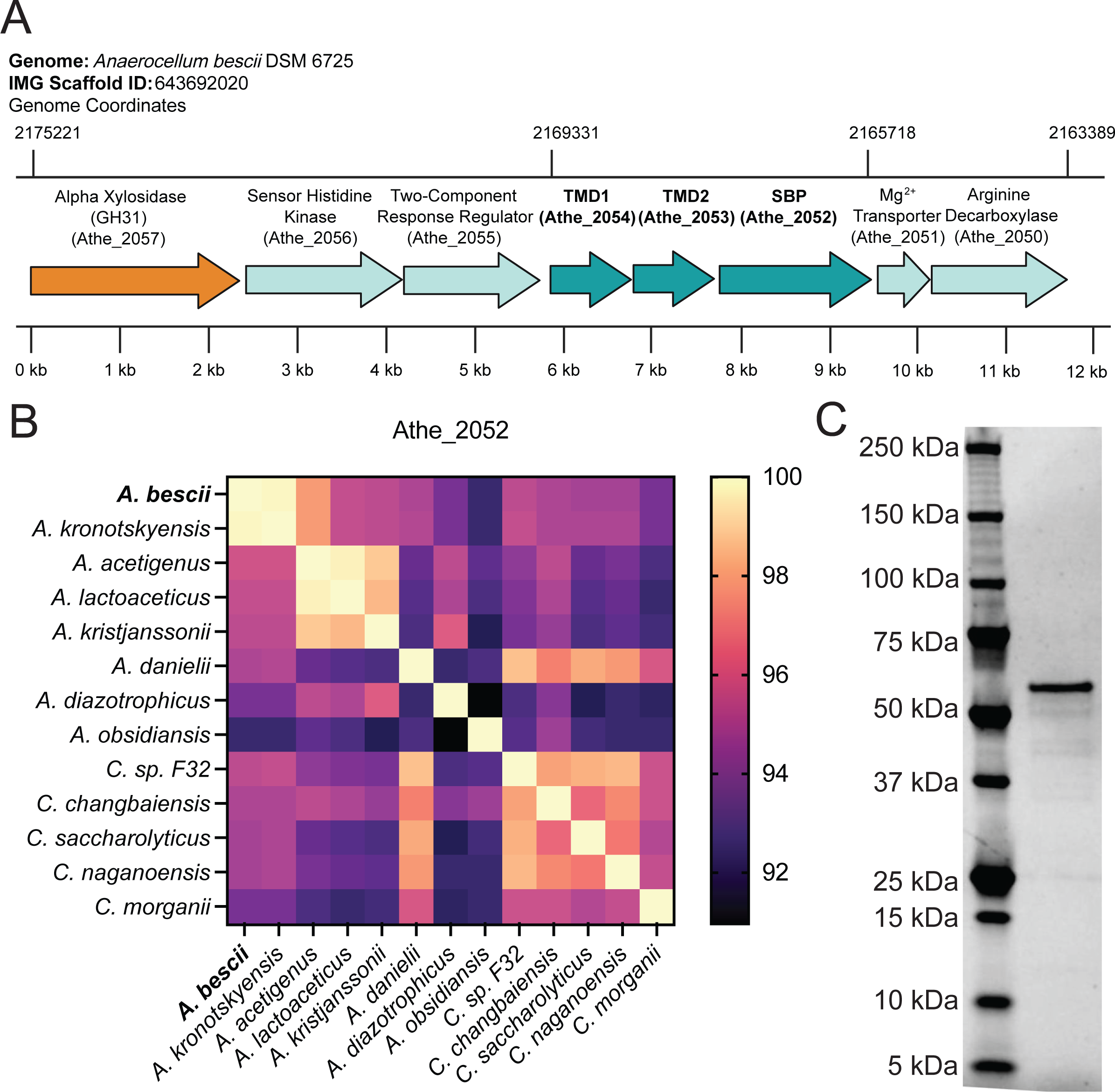
Xyloglucan ABC transporter locus and conservation across *Anaerocellum* and *Caldicellulosiruptor* species. **A)** Genomic neighborhood of the putative xyloglucan ABC transporter locus in *A. bescii* DSM6725 (IMG Genome ID: 643692002, NCBI Accession NC_012034). Coordinates are given based on IMG Scaffold ID: 643692020 and genes are identified by their locus tag (Athe_####). Genes encoding the transmembrane and substrate-binding domains of the ABC transporter complex are colored in dark cyan, while the putative α-xylosidase relevant for intracellular debranching of xyloglucan oligosaccharides is colored in orange. **B)** Protein sequence conservation of Athe_2052 across the *Anaerocellum* and *Caldicellulosiruptor* genera by BLASTp. **C)** Purified Athe_2052, without its signal peptide, run on a SDS-PAGE gel.

This xyloglucan utilization locus is broadly conserved across the *Anaerocellum* and *Caldicellulosiruptor* genera (Figure 2). While some ABC transporters such as the cello-oligosaccharide binding proteins Athe_0597 and Athe_0598 show genetic differences between the two genera, Athe_2052 is highly conserved across both (19). All *Caldicellulosiruptor* species, and all *Anaerocellum* species with the exception of *A. hydrothermalis* and *A. obsidiansis,* possess the genes encoding the extracellular SBP and two transmembrane domain components, with relatively high amino acid sequence similarity at 93.6% or greater for the substrate-binding domain. Indeed, certain *Caldicellulosiruptor* species such as *C. sp. F32, C. morganii, C. changbaiensis, C. saccharolyticus,* and *C. naganoensis* bear greater sequence similarity (with respect to the substrate-binding domain) to *A. bescii* compared to true *Anaerocellum* species *A. diazotrophicus* and *A. obsidiansis* (Figure 2).

### The Structure of Athe_2052 Reveals a Rare Structural Fold Only Found in Cluster G Substrate Binding Proteins

The three-dimensional structure of Athe_2052 was solved to a resolution of 2.43 Å (Figure 3, Table 1). The monomeric structure of Athe_2052 follows the classical Rossmann-like α/β sandwich fold of two lobes connected by a flexible hinge. The lobe containing the N-terminus, hereafter termed the N-terminal domain (NTD), is formed by a triplet of antiparallel β-sheets, surrounded by eleven α-helices and an additional pair of antiparallel β-sheets. The comparatively larger C-terminal domain (CTD) also consists of a core triplet of antiparallel β-sheets surrounded by ten α-helices, along with an additional two-stranded β-sheet near the hinge. The hinge region comprises three solvent-exposed linking regions connecting the N-terminal stretch to the C-terminal region: one loop between G423 to R451, a second between R149 to G175, and a third between Y333 to D366 (Figure 3).

**Figure 3:**
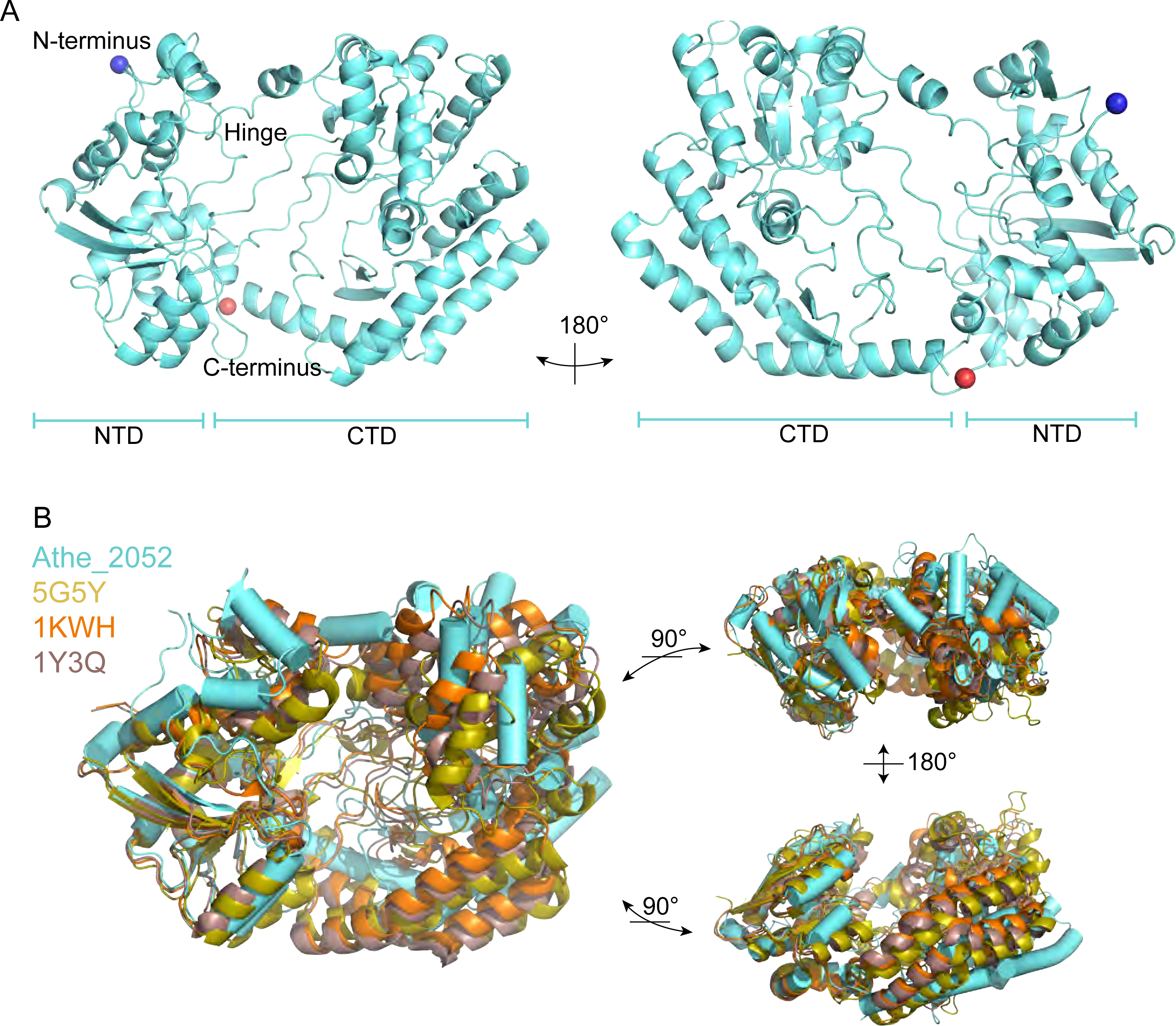
Crystal structure of Athe_2052 and comparison to cluster G SBPs. **A)** The crystal structure of Athe_2052 was solved to a resolution of 2.43 Å (PDB 9PMY). The substrate-binding protein possesses two α/β lobes, with several structural features – the N-terminus, C-terminus, and hinge region – marked on the structure. NTD = N-Terminal Domain, CTD = C-Terminal Domain. The blue and red spheres represent phosphate ions. **B)** Structural alignment of Athe_2052 with three Cluster G substrate-binding protein homologs (PDB: 5G5Y, PDB: 1KWH, PDB: 1Y3Q) obtained through a PDB DALI search illustrate similar features such as a triple polypeptide hinge region and a large ligand binding pocket.

**Table 1:** Data and refinement statistics for the apo structure of Athe_2052.

| Protein | Athe_2052 |
| --- | --- |
| PDB code | 9PMY |
| Resolution range for data (Å) | 30. - 2.43 (2.49 - 2.43) |
| Space group | P21 |
| a, b, c (Å) | 43.3, 130.0, 111.2 |
| $\alpha$, $\beta$, $\gamma$ (°) | 90.0, 93.6, 90.0 |
| Unique reflections | 46,233 (3,328) |
| Completeness (%) | 99.7 (97.4) |
| Redundancy | 4.7 (4.4) |
| $CC_{1/2}$ | 0.997 (0.679) |
| $I/\sigma(I)$ | 10.4 (1.7) |
| $R_{merge}$ | 0.095 (0.775) |
| $R_{meas}$ | 0.107 (0.880) |
| Resolution range for refinement (Å) | 29.8 - 2.43 (2.48 - 2.43) |
| Reflections used in refinement | 46,198 |
| Reflections used for R-free | 2,267 |
| $R$-work | 0.164 (0.236) |
| $R$-free | 0.218 (0.293) |
| Number of Non-Hydrogen Atoms | 8989 |
| Macromolecules | 8650 |
| Ligands/ions | 10 |
| Water | 329 |
| Protein residues | 1,066 |
| RMS (bonds) (Å) | 0.008 |
| RMS (angles) (°) | 0.915 |
| Ramachandran favored (%) | 97.1 |
| Ramachandran allowed (%) | 2.9 |
| Ramachandran outliers (%) | 0.0 |
| Rotamer outliers (%) | 2.03 |
| Clash score | 4.84 |
| Wilson B-factor (Å<sup>2</sup>) | 51.4 |
| Overall average B-factor (Å<sup>2</sup>) | 69.6 |
| Average B-factor macromolecules (Å<sup>2</sup>) | 69.9 |
| Average B-factor ligands/ions (Å<sup>2</sup>) | 75.9 |
| Average B-factor water (Å<sup>2</sup>) | 59.0 |

The three-dimensional structure of Athe_2052 reveals structural characteristics conserved across Cluster G substrate-binding proteins, as confirmed by a DALI PDB homolog search (12,13): AlgQ2 from *Sphingomonas* sp. A1 (PDB ID: 1KWH, RMSD: 4.8 Å, Z-score: 19.1), AlgQ1 from *Sphingomonas* sp. A1 (PDB: 1Y3Q, RMSD: 4.8 Å, Z-score: 20.3), FusA from *Streptococcus pneumoniae* TIGR4 (PDB: 5G5Y, RMSD: 4.7 Å, Z-score: 21.3) (20–22) (Figure 3). Firstly, Athe_2052 (56 kDa, 551 amino acids) is similar in size to AlgQ2, AlgQ1, and FusA – all of which are roughly 60 kDa. Athe_2052 possesses an expansive binding pocket volume of 5180.6 Å^3^, leaving many of its binding pocket amino acid residues exposed to the bulk solvent in the apo conformation. And like its Cluster G structural homologs, Athe_2052 possesses a flexible hinge region made up of three solvent-exposed linkers allowing for a larger range of hinge motion, and made possible by its Class II sheet topology of β_2_β_1_β_3_β_n_β_4_, which contains a domain dislocation further increasing the hinge bending range. Structural alignment of Athe_2052 also reveal a similar topology, with β-4, α-6, α-7, and β-5 secondary structures being highly conserved features of the C-terminal domain (Figure S1). However, Athe_2052 lacks a calcium-binding motif present in the other Cluster G SBPs, and of the conserved calcium-binding EF-hand motif “DPNGNG” present in other Cluster G SBPs, only “P” and the second “G” are present in Athe_2052, as part of a corresponding “HPTTDG” region (20–22).

The structural features present in Athe_2052 – and found in other Cluster G SBPs – make it well-suited to binding large substrates. Indeed, all Cluster G SBPs preferentially bind large oligosaccharides of trimeric size or greater (11–13). AlgQ1 (PDB: 1Y3Q) can bind alginate tetramers of α-L-guluronate (G) and its C5 epimer β-D-mannuronate while FusA (PDB: 5G5Y) binds fructo-oligosaccharides of tetrameric size and larger (12,13),(21,22). Outside Cluster G SBPs, DALI PDB search also revealed structural similarity between Athe_2052 and the glycosaminoglycan-binding protein from *Streptobacillus moniliformis* DSM 12112 (PDB: 5GX8, RMSD: 4.4 Å, Z-score: 21.1) (23).

### Biophysical Characterization of Athe_2052 Reveals its Binding Specificity for Xyloglucan Oligosaccharides

Differential scanning calorimetry (DSC) can be used to screen for the ligand specificity of thermophilic substrate-binding proteins (19). The presence of cognate ligands induces SBPs to adopt their more thermally stable closed conformation, corresponding to a higher melting temperature (8,13,16). DSC melting curves performed on Athe_2052 with various sugar ligands illustrate its substrate specificity (Figure 4). The native, apo-state melting temperature of Athe_2052 is 89.1 °C and increases to 102.7 °C in the presence of hydrolysis products from secretome treated xyloglucan that include the oligosaccharides xyloglucan heptasaccharide, octasaccharide, and nonasaccharide (Figure 1). As xyloglucan is made up of primarily β-1,4 linked glucans, we tested the melting temperature of Athe_2052 with various cello-oligosaccharides. In the presence of cellotriose, Athe_2052 achieves a melting temperature of 94.5 °C (Table 2). Addition of a fourth glucosyl residue in the form of cellotetraose results in an even higher melting temperature of 97.4 °C (Table 2). However, addition of these cello-oligosaccharides does not fully account for the 13.6 °C melting temperature increase observed with digested xyloglucan (Table 2). As the *A. bescii* secretome digested xyloglucan contains an abundance of xyloglucan heptasaccharide (XXXG) (Figure 1), we tested this oligosaccharide with Athe_2052, resulting in a melting temperature of 104.2 °C. The XXXG-protein mixture is the most thermostable, attaining a T_m_ exceeding even that of the xyloglucan digest, which contains XXXG among other xyloglucan derived sugars (Table 2).

**Figure 4:**
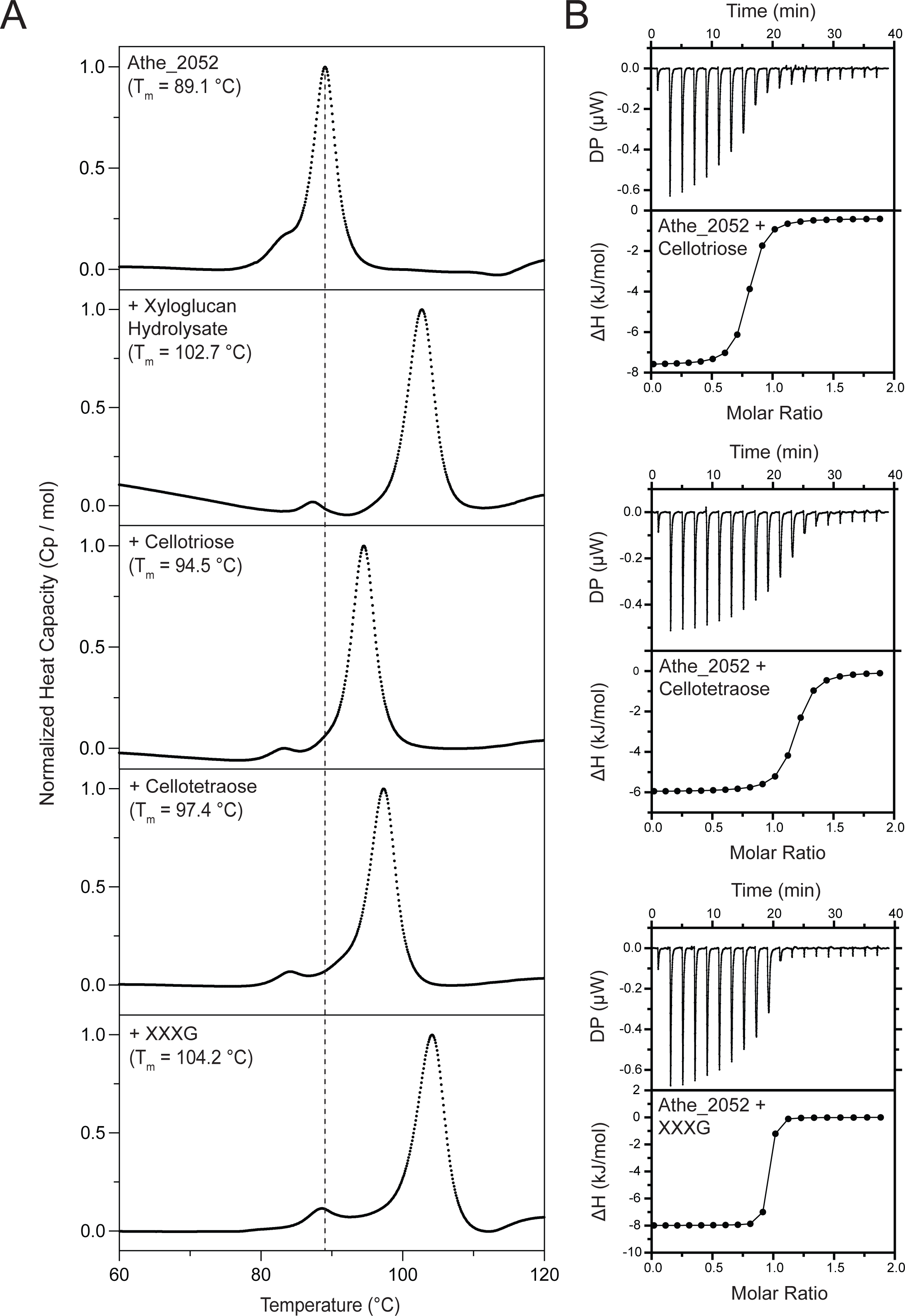
Athe_2052 ligand binding characterization by DSC and ITC. **A)** Normalized differential scanning calorimetry (DSC) screens of Athe_2052 mixed with xyloglucan hydrolysate, cellotriose, cellotetraose, and XXXG. Athe_2052 is shown to bind all tested cello-oligosaccharides and XXXG. All DSC screens were performed at a temperature range of 60 - 130°C, in 50 mM sodium acetate, 100 mM sodium chloride, 10 mM calcium chloride pH 5.5. **B)** Representative ITC screens of Athe_2052 with cellotriose, cellotetraose, and xyloglucan heptasaccharide (XXXG). Both raw isothermal titration curves and integrated binding isotherms are shown for each protein-carbohydrate mixture.

**Table 2:** Binding parameters between Athe_2052 and various sugar substrates as determined by DSC and ITC. Xyloglucan hydrolysate was not tested as a substrate via ITC due to its viscousness and chemical heterogeneity.

| Protein | Sugar | $T_m$ (°C) | $\Delta T_m$ (°C) | $n$ | $K_d$ (μM) | $K_a$ ($\times 10^5 \text{ M}^{-1}$) | $T$ (°C) | $\Delta H$ ($\frac{\text{kcal}}{\text{mol}}$) | $-T \times \Delta S$ ($\frac{\text{kcal}}{\text{mol}}$) | $\Delta G$ ($\frac{\text{kcal}}{\text{mol}}$) |
| --- | --- | --- | --- | --- | --- | --- | --- | --- | --- | --- |
| Athe_2052 | Xyloglucan hydrolysate | 102.68 | 13.61 | - | - | - | - | - | - | - |
| | Cellotriose | 94.54 | 5.47 | 0.76<br>$\pm 0.01$ | 0.247<br>$\pm 0.054$ | 40.49 | 25 | -7.24<br>$\pm 0.14$ | -1.79 | -9.02 |
| | Cellotetraose | 97.36 | 8.29 | 1.15<br>$\pm 0.01$ | 0.214 ±<br>0.029 | 46.73 | 25 | -5.93<br>$\pm 0.08$ | -3.18 | -9.10 |
| | XXXG | 104.16 | 15.09 | 0.92<br>$\pm 0.00$ | 0.014<br>$\pm 0.001$ | 714.29 | 25 | -7.99<br>$\pm 0.03$ | -2.73 | -10.70 |

To analyze the substrate-specific binding thermodynamics of Athe_2052, we measured its binding with xyloglucan heptasaccharide as well as cellotriose and cellotetraose using isothermal titration calorimetry (ITC). We obtained binding isotherms at near stoichiometric proportions in all protein-substrate combinations (Figure 4, Table 2). Confirming our DSC results, ITC measurements show that Athe_2052 possesses the highest affinity for xyloglucan heptasaccharide, with K_d_ = 14 nM.

Cellotetraose, which forms the backbone of xyloglucan heptasaccharide, is bound at a relatively low dissociation constant of K_d_ = 214 nM. Despite being one glucosyl unit shorter, trimeric cellotriose still binds with comparable binding affinity to Athe_2052 as tetrameric cellotetraose, yielding a similar dissociation constant at K_d_ = 247 nM. These K_d_ values imply a strong binding affinity and are consistent with data obtained for other known cellodextrin-specific SBPs (19,24,25). However, Athe_2052 exhibits an even stronger binding affinity for XXXG compared to undecorated glucans (K_d_ = 14 nM). The xylosyl substitutions that differentiate XXXG from undecorated β-glucans likely drive this improved binding affinity.

### Molecular Modeling Provides Structural Insights Into Recognition of Xylose-Substituted β-Glucans by Athe_2052

To understand the mechanism driving Athe_2052 selectivity for xyloglucan oligosaccharides, we investigated Athe_2052 conformational dynamics using all-atom molecular dynamics (MD) simulation of the apo, open crystal structure in the presence of its strongest binding ligand, xyloglucan heptasaccharide. A model for Athe_2052 in its holo, closed conformation was obtained from extracting the consensus pose from the MD simulation. Over the simulation, the Athe_2052 hinge bend angle decreased by 10.9°, shrinking its pocket diameter from 27.3 Å to 21.4 Å. The binding groove in Athe_2052 is the most negatively charged region of the protein, driven by an abundance of aspartic acid and glutamic acid residues (D113, D165, D167, D171, D225, D244, D245, D291, D292, D347, D449, D454, D508 and E56, E247, E487) but which becomes significantly less accessible to the bulk solvent in its holo conformation (Figure 5). Like other SBPs from *A. bescii,* Athe_2052 possesses an alkaline surface charge density (pI ∼ 9.3, Figure 5) that may make it attractive to plant cell walls, which tend to be negatively charged (26).

**Figure 5:**
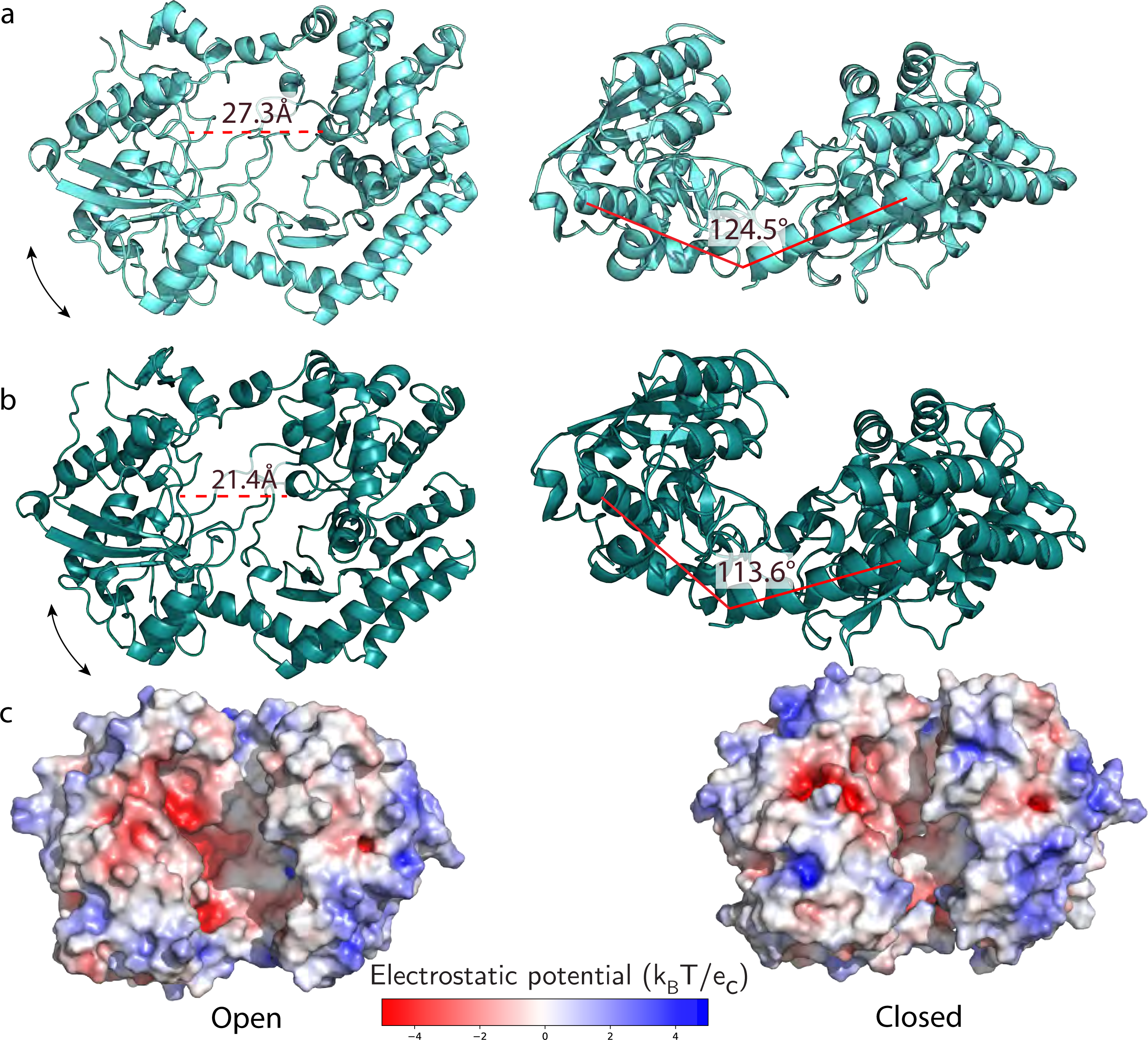
Modeling of the open and closed conformations of Athe_2052. **A)** Structural representation of Athe_2052 in its open, apo conformation and its corresponding surface charge distribution shown via a cartoon model. **B)** Molecular dynamics simulation of Athe_2052 in its closed, holo conformation, and its corresponding surface charge distribution.

We also performed docking on Athe_2052 and cellodextrin substrates. We recapitulated the same rank order of substrate preference as observed in both ITC and DSC: a top preference for XXXG, followed by secondary and tertiary affinities for cellotetraose and cellotriose, respectively (Figure 6). Corroborating ITC measurements, the free energies of binding for all three ligands are primarily driven by entropic gains due to the ejection of water molecules from the binding pocket, and second by the specific enthalpic interactions formed between the sugar and major contact residues. Interestingly, we observed a difference in the binding mechanism for XXXG and undecorated cellodextrins. As cellodextrins possess numerous hydroxyl groups that can act as both proton donors and oxygen acceptors, their binding is typically stabilized by a large network of hydrogen bonds with both protein side chains and surrounding water molecules (27). This was similarly observed for cellotriose, which forms hydrogen bonds with charged residues E56 and K111 on the non-reducing sugar ring, while aromatic residues Y109 and W311 and aliphatic residue I162 are aligned to form a hydrophobic platform along the glucan chain (Figure 6). These residues remain involved in cellotetraose binding, with the addition of W226 and N234 in forming respective van der Waals contacts and hydrogen bonds to stabilize the fourth glucosyl residue (Figure 6).

**Figure 6:**
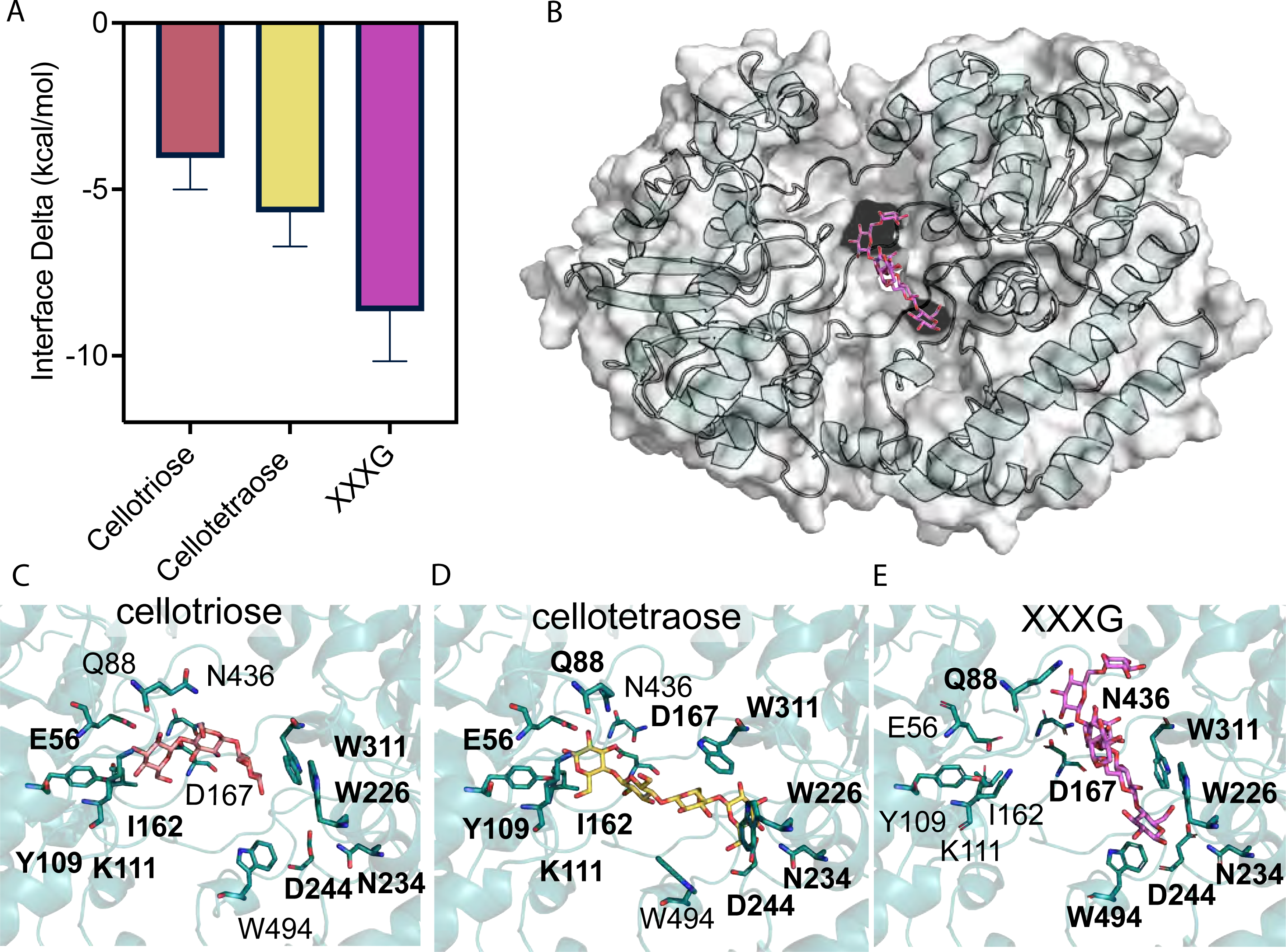
Distinct binding modes for undecorated cellodextrins versus XXXG in Athe_2052. **A)** Interface energy deltas of binding are shown for each simulation of Athe_2052 docked with experimentally tested oligosaccharides: cellotriose, cellotetraose, XXXG. **B)** Cartoon model of Athe_2052 (cyan) with its cognate substrate XXXG (purple) docked in its binding pocket. **C)** Pocket residues mediating cellotriose binding. D) Cellotetraose binding is driven by a combination of residues already involved in cellotriose binding as well as new contact residues. **E)** XXXG is bound orthogonally to ββ-glucan ligands to accommodate its α-xylose decorations.

However, XXXG appears to dock at an oblique orientation, rotated 45° relative to the pose of the undecorated cellodextrins, with a more twisted glucan conformation to accommodate its bulky side chains (Figure 6). Inspection of the binding pose for cellotetraose suggests the xylose decorations in XXXG would sterically clash with E56 and I162 (Figure 6). Consequently, a more intensive, interchain hydrogen bonding network driven by a relatively distinct set of contact residues stabilizes the longer oligosaccharide. Firstly, E56, K111, and I162 are no longer actively capping the nonreducing end of the cellodextrin backbone in XXXG; D167 and N436 instead form hydrogen bonds with the xylosyl side chains and Q88 now packs against the xylose decorations, while the three aromatic residues W226, W311, and the new addition of W494 forms a hydrophobic groove for the glucan backbone (Figure 6) (28). As was the case with cellotetraose, N234 forms a hydrogen bonding cap with the reducing end; we speculate the combination of this hydrophobic groove and cap may impose a size limit on the length of the recognizable glucan backbone (27,28). Through an unsupervised machine learning analysis, residues 161-167 (“GITTDND”), located in the hinge region between Domain I and II, were determined to be predominantly responsible for the observed open-to-closed transition. The tight packing and network of interactions around the XXXG in the closed pose likely contribute to the enthalpically driven binding observed in ITC. Though we initially surmised that bound ligands would align along the negatively charged cleft in the binding pocket, this was not the case though some overlap exists. The expanse of the binding cavity also accommodates water molecules that form a solvation shell with bound ligands (27).

## Discussion

We have shown that Gram-positive *A. bescii* naturally degrades xyloglucan into soluble oligosaccharides and possesses a conserved xyloglucan ABC transporter (Athe_2052-2054) for their uptake. We have identified Athe_2052 as the extracellular SBP that binds xyloglucan and determined its crystal structure, making it the first resolved structure of an SBP that recognizes xyloglucan. Notably, we found that Athe_2052 possesses a tertiary fold homologous to the three Cluster G SBPs that are oligosaccharide-specific but do not bind xyloglucan (20–22). This mode of xyloglucan utilization differs from TonB-dependent transport mechanisms of Gram-negative bacteria such as *B. ovatus* but parallels other Gram-positive bacteria such as *Ruminiclostridium cellulolyticum* (1,3,6).

Our crystal structure of Athe_2052 in its apo form, coupled with molecular dynamic simulations of its holo form, provide the first structural insights for the recognition of xyloglucan oligosaccharides via specific interactions with the xylosyl substitutions. These findings were quantitatively supported by biophysical experimental approaches: DSC measurements confirm that Athe_2052 achieves the greatest relative thermostability when bound to XXXG while ITC shows that XXXG is bound with a K_d_ an order of magnitude lower than those for cellotetraose or cellotriose. The obtained K_d_ = 14 nM value is comparable to those obtained for pure XXXG with a similar xyloglucan ABC transporter in *Ruminiclostridium cellulolyticum,* and is lower than dissociation constants for most ABC SBPs in lignocellulolytic bacteria (6,16,24,25).

To structurally illustrate the binding of Athe_2052 to xyloglucan oligosaccharides, we modeled the holo conformation of Athe_2052 based on the apo crystal structure and molecular dynamics simulations. Ligand docking of cellotriose and cellotetraose revealed recognition by a series of both polar residues and hydrophobic interactions, but XXXG was crucially bound in a distinct conformation owing to its bulky xylose side chains. Previously, the sub-site model has been used to describe the way extracellular SBPs from *A. bescii* can bind maltodextrins and cellodextrins of various sizes (15,16,19). However, the sub-site model is premised upon a conserved set of amino acid residues per sub-site to bind oligosaccharides of variable length but of uniform chemistry. Therefore, it is unsurprising that it cannot account for the oblique conformation XXXG adopts in its bound pose thanks to its branched side chains and the sheer size of the Athe_2052 binding pocket.

The multiple classes of proteins known to bind xyloglucans in either polymeric or oligomeric form thus far have recognized xyloglucan through its β-glucan chain. In *B. ovatus*, xyloglucan recognition is mediated by SusC-like and SusD-like cell surface glycan-binding proteins (1,3,29). These SGBPs possess glycan-binding platforms to cradle xyloglucan oligosaccharides in larger forms than what can be intracellularly imported via ABC transporters, e.g., *B. ovatus* SGBPs can bind xyloglucan tetra-desaccharides weighing at 2.5 kDa (1,3). However, crystallographic studies on these SGBPs only show xyloglucan recognition primarily through their β-glucan backbones; only a single tyrosine residue appeared to hydrophobically interact with a xylosyl residue (3). Similarly, lignocellulolytic bacteria such as *Clostridium thermocellum* possess multi-domain glycoside hydrolases known to bind to and degrade xyloglucan, but structural investigation of their Carbohydrate-Binding Module domains from family 44 (CBM44) also reveal a hydrophobic platform that tightly binds the β-glucan chain but leaves the xylosyl substitutions mostly solvent exposed (28). Thus remarkably, Athe_2052 utilizes a novel mode of oligosaccharide recognition that treats the xylosyl decorations as prominent molecular features driving ligand binding.

The scavenger lifestyle of Gram-positive, biomass degrading bacteria such as *R. cellulolyticum* and *A. bescii* may partially explain their mode of xyloglucan utilization. In nutrient-poor environments, large oligosaccharides are ideal substrates for cytoplasmic import because the cost of two ATP molecules per translocated substrate remains fixed for sugars of any size. Since the larger the oligosaccharide that is transported the greater its metabolic value, it stands to reason that XXXG should be a preferred growth substrate for *A. bescii.* Moreover, the uptake of xyloglucan oligosaccharides could be considered a “selfish uptake” strategy, whereby the oligosaccharides are transported intact and hydrolyzed into monosaccharides within the cytoplasm (30). This process prevents neighboring microbes from accessing the released sugars and has been previously reported for arabinoxylan oligosaccharide utilization in *R. cellulolyticum* (30).

The highly expressed Glucan Degradation locus (GDL; *Athe_1857-1867*) in *A. bescii* encodes secreted, multi-domain CAZymes for plant cell wall deconstruction (31). Among these, CelF (Athe_1860; GH74-CBM3-CBM3-GH48) is the key xyloglucanase. Its GH74 domain cleaves xyloglucan at unbranched glucosyl units, yielding XXXG as its primary product (32). We previously characterized the full-length CelF enzyme, demonstrating its activity on tamarind xyloglucan and its role within the *A. bescii* secretome (17,31). Here we link this extracellular activity which releases XXXG to uptake by the Athe_2052-2054 ABC transporter. Following import of xyloglucan oligosaccharides, Athe_2057, a GH31 encoded in the same locus and annotated as an α-xylosidase, is likely responsible for xylose debranching. For XXLG and XLLG, Athe_1927 (GH42, putative β-galactosidase) may play a role in removing galactosyl residues. And, while we have constrained our investigation to tamarind xyloglucan, other sources of xyloglucan can include additional fucosyl substitutions on the galactosyl residues, for which the putative α-fucosidase Athe_2072, not far from the xyloglucan ABC transporter locus, may help to remove (18).

Although CelF is a major xyloglucanase in *A. bescii*, it is expressed at relatively low levels compared to other GDL enzymes (17). This combined with its highly specialized GH74 endo-xyloglucanase function, suggests a limited role focused on xyloglucan oligosaccharide release to facilitate physical access to more abundant plant polysaccharides such as cellulose or xylan for degradation by other GDL CAZymes (17). Consistent with this, the ABC transporter genes *Athe_2052 – 2054* are generally up-regulated during growth on plant biomass, e.g., cellulose or xylan (18). However, it is also expressed at lower levels compared to more highly transcribed ABC transporters such as those for xylo-oligosaccharides (*Athe_0614 – 0616*), supporting the view that xyloglucan is not a primary carbon source but opportunistically utilized as it is released from plant biomass (18,33). We therefore propose a model that describes the mechanism of xyloglucan deconstruction and uptake in *A. bescii* (Figure 7), paralleling mechanisms described for *R. cellulolyticum* (6) and potentially generalizable across the *Anaerocellum* and *Caldicellulosiruptor* genera.

**Figure 7:**
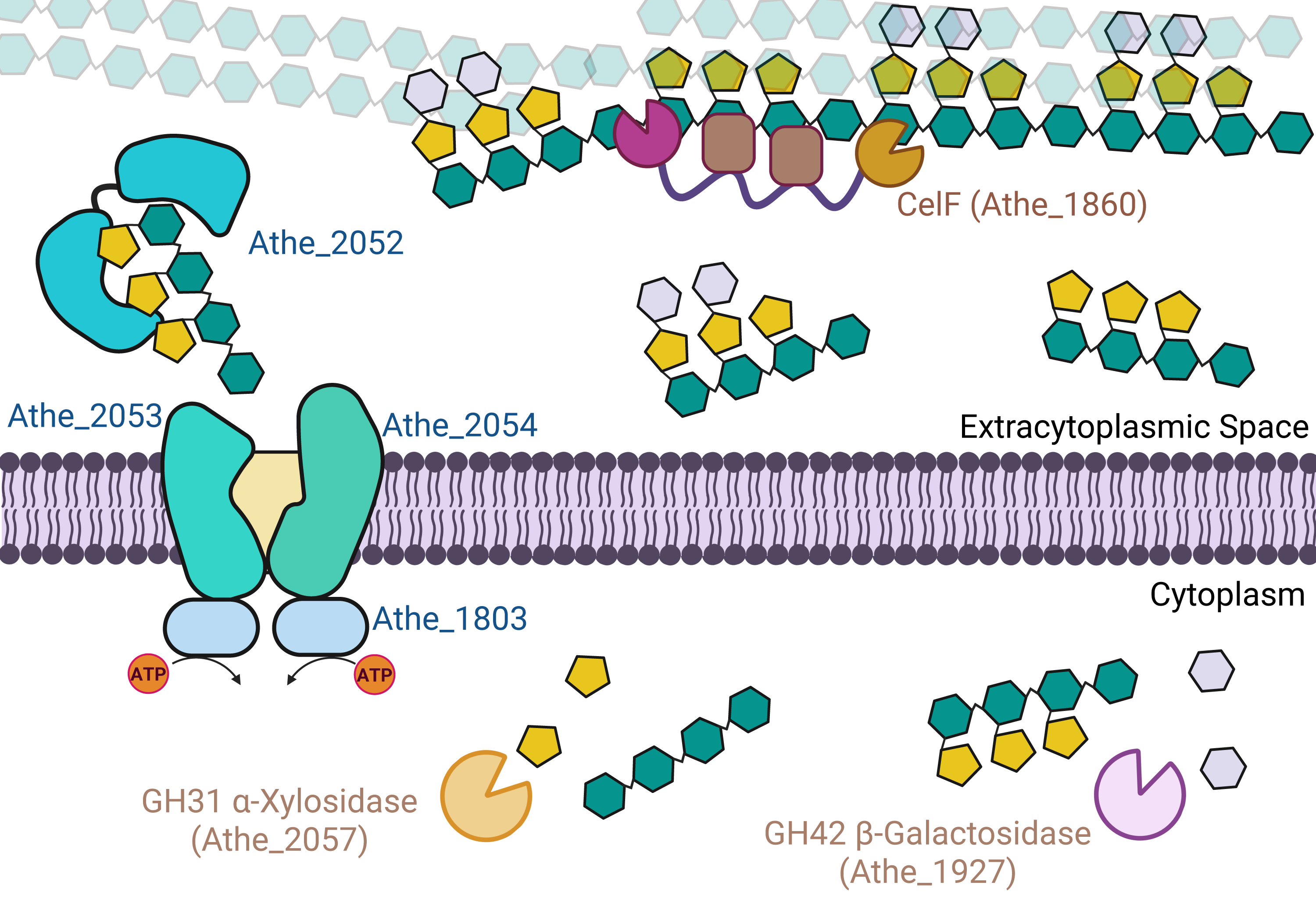
Proposed model for xyloglucan deconstruction and uptake in Gram-positive *A. bescii*. CelF (Athe_1860; GH74-CBM3-CBM3-GH48) cleaves polymeric xyloglucan to soluble oligosaccharides (primarily XXXG but also XXLG and XLLG) extracellularly. The Athe_2052–2054 ABC transporter (with ATPase Athe_1803) imports these oligosaccharides intact. In the cytoplasm, Athe_2057 (α-xylosidase; GH31) and Athe_1927 (β-galactosidase; GH42) debranch side chains, releasing monosaccharides and undecorated β-glucans for catabolism.

It has been recently discovered that Gram-positive bacteria utilize xyloglucan in a manner distinct from Gram-negative bacteria. Instead of component monosaccharides, soluble xyloglucan oligosaccharides are imported into the cytoplasm via ATP-coupled hydrolysis (1,6). Guided by our crystal structure of Athe_2052, we have illustrated the hitherto unknown structural basis for xyloglucan oligosaccharide recognition in a Gram-positive bacterium, where tight sidechain packing around XXXG drives high binding affinity. Indeed, the nature of this substrate specificity is relatively uncommon: other known xyloglucan-binding proteins (CBM6s and SGBPs) only strongly bind the β-glucan backbone with minimal side chain recognition, while individual SBPs typically only recognize a specific chemistry (3,15,16,24,28,34). We also show that the xyloglucan-binding SBP Athe_2052 in *A. bescii* is structurally unusual, and belongs to a distinct cluster of large SBPs that bind complex carbohydrate substrates. Altogether, this work advances fundamental understanding of how this important plant biomass substrate is imported by bacteria, with broad significance in microbial carbohydrate utilization across various ecological niches.

## Materials and Methods

### Bacteria & Growth Conditions

Strains of *E. coli,* including NEB 5-ɑ (New England Biolabs) and BL21(DE3) pRosetta2 (EMD Millipore), were routinely plated on Luria-Bertani agar (10 g/L tryptone, 5 g/L yeast extract, 10 g/L NaCl, plus 1.5% agar). All plates contained 50 μg/mL kanamycin for selection. *E. coli* strains were all cultured in Luria-Bertani broth supplemented with an additional 24 g/L yeast extract. For BL21(DE3) pRosetta2 strains, both plates and cultures were supplemented with 33 μg/mL chloramphenicol. *A. bescii* DSMZ 6725, provided by the lab of Dr. Robert Kelly (Department of Chemical & Biomolecular Engineering, North Carolina State University), was cultured on the complex medium C516 (5 g/L maltose, 0.5 g/L yeast extract, and 40 μM uracil) (35,36). Late exponential-phase *A. bescii* cells were harvested by centrifugation at 5,000 x g for 20 min. Genomic DNA was isolated from the resulting cell pellet using a Quick-DNA Miniprep kit (Zymo Research) and quantified via a UV-Vis Spectrophotometer Nanodrop (Thermo Scientific).

### Chemicals

The following monosaccharides and oligosaccharides were used in this study: D-Xylose (> 99%, Thermo Scientific), D-Glucose (Thermo Scientific), D-Cellobiose (> 98%, Acros Organics), D-Cellotriose (> 95%, Neogen Corporation), D-Cellotetraose (> 90%, Neogen Corporation), Xyloglucan Heptasaccharide (Biosynth). The polysaccharide xyloglucan from tamarind seed (reference code P-XYGLN) was also used (Neogen).

### Substrate Binding Protein Cloning

Athe_2052 cloning and production methods closely follow methods used previously for other *A. bescii* SBPs described in Tjo et al. (16). Briefly, the gene encoding the putative xyloglucan-binding protein Athe_2052 without its signal peptide as predicted by SignalP 5.0 was PCR-amplified from *A. bescii* DSMZ 6725 genomic DNA (37). PCR was conducted with Phusion polymerase (New England Biolabs) and primers listed in Table S1, with adherence to manufacturer protocol. Athe_2052 was inserted into a pRSF1-b backbone (courtesy of the Kelly Lab at North Carolina State University). Gene insertion utilized Gibson Assembly with the NEB HiFi Assembly Mastermix, according to manufacturer instructions. An N-terminal hexa-histidine tag preceded the gene encoding Athe_2052 to enable purification by immobilized metal affinity chromatography (IMAC). The resulting plasmid was designated pHT006. This vector was introduced into *E. coli* NEB 5-ɑ cells under 50 μg/mL kanamycin selection. The plasmid was prepared and purified via a Zymo Research Plasmid Miniprep - Classic kit (Zymo Research) and verified using Plasmidsaurus’ whole-plasmid sequencing service. The sequence-confirmed pHT006 plasmid was transformed into *E. coli* BL21(DE3) pRosetta2 (EMD Millipore) under dual selection pressures of 50 μg/mL kanamycin and 33 μg/mL chloramphenicol.

### Protein Expression & Purification

The *E. coli* BL21(DE3) pRosetta2 strain bearing the pHT006 vector was cultured in 1 L of ZYM-5052 auto-induction medium supplemented with 50 μg/mL kanamycin and 33 μg/mL chloramphenicol (38). Cultures were grown overnight in 2.8 L shake flasks at 37 °C and shake speed of 250 rpm. After 20 hours of growth, cells were harvested by centrifugation at 4,500 x g for 20 min. Cell pellets were stored at -20 °C. To resuspend cells, IMAC Buffer A (20 mM sodium phosphate monobasic, 500 mM sodium chloride, pH 7.4) was added at a ratio of 10 mL buffer per 1 gram of wet pellet. Pellets were lysed using an Emulsiflex-C5 High-Pressure Homogenizer (Avestin) with house air set to 30 psi. The lysate was subjected to a pressure of ∼100,000 kPa per pass; three passes were performed for maximal lysis. The resultant lysate was heat treated at 68 °C for 30 min, followed by centrifugation at 36,000 × g for 30 min. The collected supernatant was filtered through a 0.2 μm PES filter and subsequently loaded onto a 5 mL HisTrap HP Nickel-Sepharose column (Cytiva) using a BioRad NGC 10 FPLC system (Bio-Rad). Purified protein was analyzed on 4–20% Mini-PROTEAN® TGX Stain-Free protein gels (Bio-Rad), using the Precision Plus Protein Standard (Bio-Rad) as the molecular weight marker. Elution fractions with high protein purity and yield were pooled, concentrated, and exchanged into *A. bescii* secretome buffer (50 mM Sodium Acetate, 100 mM Sodium Chloride, 10 mM Calcium Chloride pH 5.5) using a 10 kDa MWCO PES 20 mL Spin-X Concentrator (Corning) at a speed of 3,600 x g. A bicinchoninic acid (BCA) assay (Thermo Fisher Scientific) was used to verify concentration of pHT006 protein to 5 mg/mL.

### Xyloglucan Hydrolysis

1 L of *A. bescii* culture was grown in base complex media C516 (0.5 g/L yeast extract, 40 μM uracil) supplemented with 2.5 g/L of xylose, 2.5 g/L of glucose, and 0.5 g/L of tamarind seed xyloglucan to maximize expression of GHs active on xyloglucan. After reaching stationary phase at 20 hours of growth, *A. bescii* culture was pelleted using 50 mL Falcon Conical Centrifuge Tubes (Corning) via centrifugation at 6,000 x g after which the supernatant was collected. The cell supernatant was concentrated 200-fold to 5 mL total volume via centrifugation at 3,600 x g using a 10 kDa MWCO PES 20 mL Spin-X Concentrator (Corning) and buffer exchanged into *A. bescii* secretome buffer (50 mM sodium acetate, 100 mM sodium chloride, 10 mM calcium chloride pH 5.5). An A280 nm measurement on the Nanodrop was used to verify secretome protein content diluted to 1 mg/mL. 5 g/L of xyloglucan was hydrolyzed in a total reaction volume of 4800 μL with a secretome concentration of 0.2 mg/mL at a temperature of 70 °C for 6 hours. Raw xyloglucan hydrolysate was centrifuged using a 10 kDa MWCO PES 20 mL Spin-X Concentrator (Corning) at a speed of 3,500 x g and temperature of 4 °C to separate exoproteomic content in the retentate from soluble sugars in the flow-through. The flow-through was collected and its released sugar content analyzed via the 3,5-dinitrosalicyclic acid (DNS) reducing sugar assay in 96-well format (39). Each well contained thirty-six microliters of hydrolysis reaction reacted with DNS at a 1:2 ratio, then diluted with 160 μ*L* of Milli-Q water and read at 540 nm using a BioTek Synergy H1 microplate-reader. A standard curve was constructed using D-xylose standards to determine reducing sugar equivalents per unit absorbance, and used to determine the reducing sugar concentration of the xyloglucan hydrolysate.

### Liquid Chromatography - Mass Spectrometry (LC-MS)

LC-MS analysis was conducted using an Agilent 6530 QTOF with Agilent 1260 LC operating system. Xyloglucan hydrolysate sample was flowed through an Agilent HI-419 PLEX Na (Octo) 300 x 7.7 mm column heated to 80 ℃, followed by electrospray ionization in positive ion mode to obtain the mass spectra of hydrolysate contents. The sugar sample was run using ultra-pure Milli-Q water as mobile phase, at a flow rate of 0.5 mL/min and a total run time of 30 minutes. Agilent MassHunter software was used to extract ion chromatograms and mass spectra for xyloglucan oligosaccharides, reported as [M+Na]^+^ adducts.

### Crystallization and Data Collection

Purified Athe_2052 at a concentration of 5 mg/mL was crystallized via sitting drop vapor diffusion. Each drop contained 1.5 μL of Athe_2052, buffered in 50 mM Tris pH 7.0, 150 mM NaCl, mixed with 1.5 μL of a solution containing 46 mM potassium phosphate (monobasic) and PEG 8,000 (25% w/v). Diffraction quality crystals in the form of thin needles grew within approximately one week at a temperature of 25 °C.

Data were collected at beam line 17-ID-1 (AMX) at Brookhaven National Lab, Upton, New York, USA. Data were indexed and integrated using XDS and scaled using AIMLESS to a maximum resolution of 2.43 Å (Table 1) (40,41). Crystals grew in space group *P2_1_* with cell dimensions a=43.3 Å b=130.0 Å c=111.2 Å and β=93.6° with two molecules in the asymmetric unit. The structure was determined by molecular replacement using the program PHASER starting from the AlphaFold 2.0 model of Athe_2052 and iteratively rebuilt with COOT and refined with PHENIX.REFINE at a resolution of 2.43 Å (42–44). The final model of Athe_2052 is dimeric and contains two phosphate (PO_4_) groups. Three hundred and twenty-nine water molecules are also present in the model. Final model statistics are given in Table 1. The structure has been deposited in the Protein Data Bank (PDB) with code 9PMY (45).

### Differential Scanning Calorimetry (DSC)

Biophysical measurements on Athe_2052 follow methods used previously in Tjo et al. 2025 (16). Briefly, melting curves of the hexa-histidine-tagged Athe_2052 protein were analyzed via DSC on the MicroCal PEAQ-DSC instrument (Malvern Panalytical) at the Princeton University Biophysics Core Facility. The protein was tested against a mixture of *A. bescii* secretome-digested xyloglucan polysaccharide, as well as a series of pure oligosaccharides – cellotriose, cellotetraose, xyloglucan heptasaccharide. Each sample of protein and pure sugar consisted of 2 mg/mL protein and 5 mM sugar in the same *A. bescii* secretome buffer (50 mM sodium acetate, 100 mM sodium chloride, 10 mM calcium chloride pH 5.5), which was also used as a reference. Xyloglucan hydrolysate sample was diluted to a reducing sugar equivalent of 2 mg/mL, and correspondingly mixed with 2 mg/mL of protein. The instrument was run at a scan rate of 3 °C / min, reaching a maximum temperature of 130 °C. Raw data comprising the heat capacity (C_p_) was plotted against temperature and subsequently normalized by each run’s maximum heat capacity, C_p, max_. This maximum value is defined as C_p_ (T = T_m_), where T_m_ indicates the melting temperature of the protein mixture. Each protein-sugar mixture was assigned a ΔT_m_ value given by ΔT_m_ = T_m, holo_ – T_m, apo_, where T_m, holo_ denotes the melting temperature of the protein-sugar combination and T_m,apo_ denotes the melting temperature of the protein without sugar.

### Isothermal Titration Calorimetry (ITC)

Similar methods are performed as described in Tjo et al. 2025 (16). MicroCal PEAQ Isothermal Titration Calorimeter (Malvern Panalytical) from the Princeton University Biophysics Core Facility was used to perform all measurements, at a temperature of 25 °C. Titrations were performed using a protein sample volume of 280 μL at 50 μM concentration in the device cell, and carbohydrate ligand volume of 40 μL and 500 μM in the syringe. Cell stir speed was set to 750 rpm due to protein viscosity. Protein in the cell was injected with a primer of 0.2 μL ligand, followed by 19 consecutive injections at 2.0 μL volume each and 120 s intervals in between injections. Non-linear regression based on single site binding model was used to determine integrated heat effects (Microcal PEAQ-ITC Analysis). Thermodynamic and binding parameters were obtained from the fitted isotherms using the Gibbs Free Energy form ΔG = -RTln(K_d_). The slope of the binding isotherm at the equivalence point was used to obtain the dissociation constant K_d_. Its inverse, K_a_, is defined as the association rate constant. 280 μL Milli-Q water was used in the reference cell.

### Molecular Simulation

Starting from the apo, open crystal structure obtained in this paper (PDB: 9PMY), xyloglucan heptasaccharide (CID 44630346, National Center for Biotechnology Information, PubChem Compound Summary) was positioned in the central cavity using AutoDock Vina (46). The protein and ligand structure was prepared for simulation using Solution Builder in CHARMM-GUI (47,48). The system was solvated in a cubic box of length 107 nm, where the protein was at least 1 nm away from any box edge. The parameters used for the protein are Amber ff19SB (49); the parameters used for the sugar are Amber GLYCAM-06j (50); water molecules were parameterized with TIP4P-D; the system was neutralized with NaCl to a total ion concentration of 150 mM.

Molecular dynamics simulations were performed in Gromacs v. 2022.5 patched with Plumed v.2.8.3. To minimize the energy of the initial structure, 5000 steps of steepest descent minimization were performed. To equilibrate the temperature of the system, 125 ps of NVT simulation at 303K were performed using the velocity-rescaling thermostat (τ_T = 1 ps), where the protein and ligand were coupled separately from the water and ions in the system. 1 ns of NPT equilibration was performed using the stochastic cell rescaling barostat at 1 bar (τ_T = 1 ps, τ_P = 5 ps). 50 ns of production simulation were performed, also in the NPT ensemble at 1 bar and 303 K. Hydrogen atoms were constrained using LINCS (51); electrostatics were computed with particle-mesh Ewald summation (52), using a Coulombic cutoff of 0.9 nm. Trajectory analysis was done using the Python package MDTraj (53) and the trajectory was visualized using Visual Molecular Dynamics (VMD) (54).

A consensus closed, holo pose was determined from k-means clustering on principal component analysis (PCA) from dihedral angles over the course of the trajectory. The ligand was deleted from this pose to generate a closed, apo pose that served as the input pose for docking cellotriose and cellotetraose. Initial bound cellodextrin poses were determined using AutoDock Vina, then flexible induced-fit docking was performed in Rosetta using RosettaLigand through the ROSIE server (55–57). Ligand conformers were created using BCL (58); 200 independent ligand/protein structures were produced, and the lowest energy pose for each ligand was evaluated structurally. KDEEP was used to assess the interface energy of the bound cellotriose, cellotetraose, and xyloglucan heptasaccharide (59). Pocket volume measurements were performed with CASTpFold. All protein structure figures were prepared using PyMOL (60).

### Bioinformatics

Four close homologs to Athe_2052 were identified using a PDB DALI sequence similarity search (61). These homologs include AlgQ2 from *Sphingomonas* sp. A1 (PDB ID: 1KWH, RMSD: 4.8 Å, Z-score: 19.1), AlgQ1 from *Sphingomonas* sp. A1 (PDB: 1Y3Q, RMSD: 4.8 Å, Z-score: 20.3), FusA from *Streptococcus pneumoniae* TIGR4 (PDB: 5G5Y, RMSD: 4.7 Å, Z-score: 21.3), and the glycosaminoglycan-binding protein from *Streptobacillus moniliformis* DSM 12112 (PDB: 5GX8, RMSD: 4.4 Å, Z-score: 21.1) (20–23). Structure-based sequence alignment to identify similarities in secondary and tertiary protein structure, between Athe_2052 and its homologs, was performed using PROMALS3D and ESPript 3.0 visualization software (62,63). The solved crystal structure of Athe_2052 (PDB: 9PMY from this work) was used to assign a secondary protein structure to the alignment.

## Data Availability and Supporting Information

The protein crystal structure for apo Athe_2052 has been deposited in the Protein Data Bank with the accession number 9PMY. Supporting information accompanies this article. All other data can be found in the manuscript and supplementary information.

## Acknowledgments

This research used resources of the National Synchrotron Light Source II, a U.S. Department of Energy 475 (DOE) Office of Science User Facility operated for the DOE Office of Science by Brookhaven National Laboratory under Contract No. DE-SC0012704. The Center for BioMolecular Structure (CBMS) is primarily supported by the National Institutes of Health, National Institute of General Medical Sciences (NIGMS) through a Center Core P30 Grant (P30GM133893), and by the DOE Office of Biological and Environmental Research (KP1605010). Molecular dynamics simulations were performed using the Princeton Research Computing resources at Princeton University. Mass spectrometry data were collected on an instrument purchased with a supplement to NIH grant GM107036 to A.J.L.

## Author Contributions

**Hansen Tjo**: Conceptualization, Data curation, Formal analysis, Investigation, Visualization, Writing – Original draft, Writing – Review & Editing.

**Virginia Jiang**: Conceptualization, Data curation, Formal analysis, Investigation, Visualization, Writing – Original draft, Writing – Review & Editing.

**Philip D. Jeffrey**: Data curation, Formal analysis, Investigation, Writing – Review & Editing.

**Angela Zhu**: Data curation, Formal analysis, Investigation, Visualization, Writing – Review & Editing.

**A. James Link**: Funding acquisition, Resources, Supervision, Writing – Review & Editing.

**Jerelle A. Joseph**: Funding acquisition, Resources, Supervision, Writing – Review & Editing.

**Jonathan M. Conway**: Conceptualization, Funding acquisition, Project administration, Resources, Supervision, Writing – Review & Editing.

