## Supplementary Information for "Structural insights into xyloglucan recognition by an ABC transporter from a Gram-positive, thermophilic bacterium"

Author Affiliations:

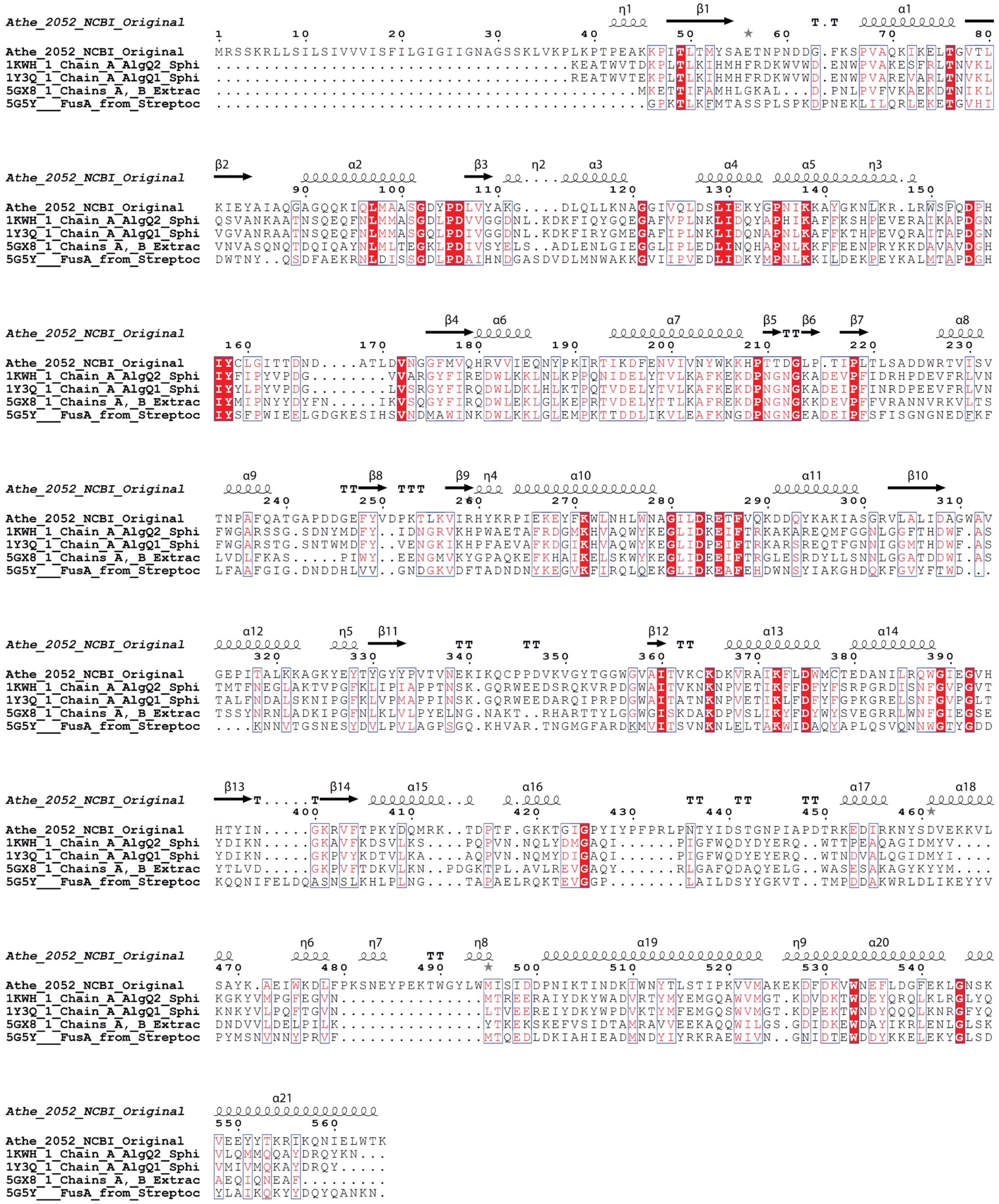

**Figure S1:** Structure-based amino acid sequence alignment of Athe_2052 (with signal peptide included) with its closest DALI homologs: AlgQ2 from *Sphingomonas* sp. A1 (PDB ID: 1KWH, RMSD: 4.8 Å, Z-score: 19.1), AlgQ1 from *Sphingomonas* sp. A1 (PDB ID: 1Y3Q, RMSD: 4.8 Å, Z-score: 20.3), FusA from *Streptococcus pneumoniae* TIGR4 (PDB ID: 5G5Y, RMSD: 4.7 Å, Z-score: 21.3), and the glycosaminoglycan-binding protein from *Streptobacillus moniliformis* DSM 12112 (PDB: 5GX8, RMSD: 4.4 Å, Z-score: 21.1). Residues shaded in red indicate full residue conservation between Athe_2052 and all four homologs. Boxes are drawn wherever aligned residues are chemically similar for at least four out of the five proteins (sequence consensus with a threshold of ≥80%).

**Table S1:** Amino acid sequences of all proteins analyzed in this study. Residues comprising the signal peptide sequence are marked in red. Only sequences obtained from NCBI contain signal peptides.

| Protein | Amino Acid Sequence |
|---|---|
| Athe_2052<br>(NCBI accession:<br>WP_015908408.1) | MRSSKRLLSILSIVVVISFILGIGIIGNAGSSKLVKPLKPTPEAKKPITLTMY<br>SAETNPNDDGFKSPVAQKIKELTGVTLKIEYAIAQGAGQQKIQLMAASGDYPD<br>LVYAKGDLQLLKNAGGIVQLDSLIEKYGPNIKKAYGKNLKRLRWSPQDPHIYC<br>LGITTDNDATLDVNGGFMVQHRVVIEQNYPKIRTIKDFENVIVNYWKKHPTTD<br>GLPTIPLTLSADDWRTVISVTNPAFQATGAPDDGEFYVDPKTLKVIRHYKRPI<br>EKEYFKWLNHLWNAGILDRETFVQKDDQYKAKIASGRVLALIDAGWAVGEPIT<br>ALKKAGKYEYTYGYYPVTVNEKIKQCPPDVKVGYTGGWGVAITVKCKDKVRAI<br>KFLDWMCTEDANILRQWGIEGVHHTYINGKRVFTPKYDQMRKTDPTFGKKTGI<br>GPYIYPFPRLPNTYIDSTGNPIAPDTRKEDIRKNYSDVEKKVLSAYKAEIWKD<br>LFPKSNEYPEKTWGYLWMISIDDPNIKTINDKIWNYTLSTIPKVVMAKEKDFD<br>KVWNEFLDGFEKLGNSKVEEYYTKRIKQNIELWTK |
| Athe_2052<br>(Purified with His-tag) | MAHHHHHHVDDDDKGSSKLVKPLKPTPEAKKPITLTMYSAETNPNDDGFKSPV<br>AQKIKELTGVTLKIEYAIAQGAGQQKIQLMAASGDYPDLVYAKGDLQLLKNAG<br>GIVQLDSLIEKYGPNIKKAYGKNLKRLRWSPQDPHIYCLGITTDNDATLDVNG<br>GFMVQHRVVIEQNYPKIRTIKDFENVIVNYWKKHPTTDGLPTIPLTLSADDWR<br>TVISVTNPAFQATGAPDDGEFYVDPKTLKVIRHYKRPIEKEYFKWLNHLWNAG<br>ILDRETFVQKDDQYKAKIASGRVLALIDAGWAVGEPITALKKAGKYEYTYGYY<br>PVTVNEKIKQCPPDVKVGYTGGWGVAITVKCKDKVRAIKFLDWMCTEDANILR<br>QWGIEGVHHTYINGKRVFTPKYDQMRKTDPTFGKKTGIGPYIYPFPRLPNTYI<br>DSTGNPIAPDTRKEDIRKNYSDVEKKVLSAYKAEIWKDLFPKSNEYPEKTWGY<br>LWMISIDDPNIKTINDKIWNYTLSTIPKVVMAKEKDFDKVWNEFLDGFEKLGN<br>SKVEEYYTKRIKQNIELWTK |
| PDB: 5G5Y | GPKTLKFMTASSPLSPKDPNEKLILQRLEKETGVHIDWTNYQSDFAEKRNLDI<br>SSGDLPDAIHNDGASDVDLMNWAKKGVIIPVEDLIDKYMPNLKKILDEKPEYK<br>ALMTAPDGHIYSFPWIEELGDGKESIHSVNDMAWINKDWLKKLGLEMPKTTDD<br>LIKVLEAFKNGDPNGNGEADEIPFSFISGNGNEDFKFLFAAFGIGDNDDHLVV<br>GNDGKVDFTADNDNYKEGVKFIRQLQEKGLIDKEAFEHDWNSYIAKGHDQKFG<br>VYFTWDKNNVTGSNESYDVLPVLAGPSGQKHVARTNGMGFARDKMVITSVNKN<br>LELTAKWIDAQYAPLQSVQNNWGTYGDDKQQNIFELDQASNSLKHLPLNGTAP<br>AELRQKTEVGGPLAILDSYYGKVTTMPDDAKWRLDLIKEYYVPYMSNVNNYPR<br>VFMTQEDLDKIAHIEADMNDYIYRKRAEWIVNGNIDTEWDDYKKELEKYGLSD<br>YLAIKQKYYDQYQANKN |
| PDB: 1KWH | KEATWVTDKPLTLKIHMHFRDKWVWDENWPVAKESFRLTNVKLQSVANKAATN<br>SQEQFNLMMASGDLPDVVGGDNLKDKFIQYGQEGAFVPLNKLIDQYAPHIKAF<br>FKSHPEVERAIKAPDGNIYFIPYVPDGVVARGYFIREDWLKKLNLKPPQNIDE<br>LYTVLKAFKEKDPNGNGKADEVPFIDRHPDEVFRLVNFWGARSSGSDNYMDFY<br>IDNGRVKHPWAETAFRDGMKHVAQWYKEGLIDKEIFTRKAKAREQMFGGNLGG<br>FTHDWFASTMTFNEGLAKTVPGFKLIPIAPPTNSKGQRWEEDSRQKVRPDGWA<br>ITVKNKNPVETIKFFDFYFSRPGRDISNFGVPGVTYDIKNGKAVFKDSVLKSP<br>QPVNNQLYDMGAQIPIGFWQDYDYERQWTTPEAQAGIDMYVKGKYVMPGFEGV<br>NMTREERAIYDKYWADVRTYMYEMGQAWVMGTKDVDKTWDEYQRQLKLRGLYQ<br>VLQMMQQAYDRQYKN |
| PDB: 5GX8 | MKETTIFAMHLGKALDPNLPVFVKAEKDTNIKLVNVASQNQTDQIQAYNLMLT<br>EGKLPDIVSYELSADLENLGIEGGLIPLEDLINQHAPNLKKFFEENPRYKKDA<br>VAVDGHIYMIPNYYDYFNIKVSQGYFIRQDWLEKLGLKEPRTVDELYTTLKAF |

| | |
|---|---|
| | REKDPNGNGKKDEVPFFVRANNVRKVLTSLVDLFKASPIWYEENGMVKYGPAQ<br>KEFKHAIKELSKWYKEGLIDEEIFTRGLESRDYLLSNNLGGATDDWIASTSSY<br>NRNLADKIPGFNLKLVLPYELNGNAKTRHARTTYLGGWGISKDAKDPVSLIKY<br>FDYWYSVEGRRLWNFGIEGSEYTLVDGKPVFTDKVLKNPDGKTPLAVLREVGA<br>QYRLGAFQDAQYELGWASESAKAGYKYYMDNDVVLDELPILKYTKEKSKEFVS<br>IDTAMRAVVEEKAQQWILGSGDIDKEWDAYIKRLENLGLSKAEQIQNEAF |
| PDB: 1Y3Q | REATWVTEKPLTLKIHMHFRDKWVWDENWPVAREVARLTNVKLVGVANRAATN<br>SQEQFNLMMASGQLPDIVGGDNLKDKFIRYGMEGAFIPLNKLIDQNAPNLKAF<br>FKTHPEVQRAITAPDGNIYYLPYVPDGLVSRGYFIRQDWLDKLHLKTPQTVDE<br>LYTVLKAFKEKDPNGNGKADEIPFINRDPEEVFRLVNFWGARSTGSNTWMDFY<br>VENGKIKHPFAEVAFKDGIKHVAQWYKEGLIDPEIFTRKARSREQTFGNNIGG<br>MTHDWFASTALFNDALSKNIPGFKLVPMAPPINSKGQRWEEDARQIPRPDGWA<br>ITATNKNPVETIKLFDFYFGPKGRELSNFGVPGLTYDIKNGKPVYKDTVLKAA<br>QPVNNQMYDIGAQIPIGFWQDYEYERQWTNDVALQGIDMYIKNKYVLPQFTGV<br>NLTVEEREIYDKYWPDVKTYMFEMGQSWVMGTKDPEKTWNDYQQQLKNRGFYQ<br>VMIVMQKAYDRQY |

**Table S2:** Primers used in this study.

| Primer | Sequence (5' – 3') | Application |
|---|---|---|
| HT001_F | GCTGCCACCGCTGAGCAATAACTAG | pRGB001 Vector.FOR |
| HT002_F | CTTGTCGTCGTCATCCACGTGATG | pRGB001 Vector.REV |
| HT013 | CCATCACGTGGATGACGACGACAAGGGAAGTTCAAAGCTTGTGAAGCC | Athe_2052 FOR Primer |
| HT014 | AGTTATTGCTCAGCGGTGGCAGCTTATTTTGTCCACAATTCAATGTTTT GCTTGATTC | Athe_2052 REV Primer |

**Table S3:** Docking results for cello-oligosaccharide substrates and xyloglucan heptasaccharide with Athe_2052. Each protein-ligand combination was simulated two hundred times (n=200).

| | Athe_2052 | |
|---|---|---|
| Ligand | $\Delta G$ (kcal/mol) | Standard Deviation (kcal/mol) |
| Cellotriose | -4.0277 | 0.9767 |
| Cellotetraose | -5.6625 | 1.0557 |
| Xyloglucan Heptasaccharide (XXXG) | -8.6390 | 1.5272 |
